# Evaluation of degron motifs in *Escherichia coli* using a fluorescent reporter

**DOI:** 10.64898/2026.03.07.710301

**Authors:** Matylda Anna Izert-Nowakowska, Partycja Emanuela Szybowska, Maria Magdalena Klimecka, Maria Wiktoria Górna

**Affiliations:** Structural Biology Group, Biological and Chemical Research Centre, Faculty of Chemistry, University of Warsaw, Warsaw, Poland

## Abstract

Fluorescent reporters provide a useful tool for studying degron motifs. Fusing a degron of interest to a fluorescent protein allows to accurately track protein levels overtime to characterise the degradation kinetics of studied degrons. Here we describe a rapid and simple method to study degron peptides in *Escherichia coli* using plasmid-encoded eGFP-degron fusion constructs. The described methods provide an accessible workflow to evaluate degrons. We provide protocols for generation of pBAD plasmids encoding the studied constructs and two different methods for evaluating degrons – an end-point fluorescence measurement on agar plates and a kinetic measurement in liquid cultures in a 96-well format for high-throughput degron studies.

## 1. Introduction

Degron motifs in bacteria determine protein turnover by regulating protein stability [1]. They drive protein degradation allowing bacteria to maintain the correct expression levels of essential proteins in response to external stimuli. This allows the bacteria to quickly respond to environmental stress and adapt to the changing conditions. Fluorescent reporter assays are a powerful tool to study peptide degrons and their ability to drive protein degradation [2]. Isolating a degron motif and fusing it to a non-native fluorescent protein allows for studying specific degrons in their cellular context. This can be beneficial for identification of new degradation motifs or evaluating engineered degrons for their use in synthetic applications [3, 4]. Here we present a rapid and simple screening method using plasmid-encoded fluorescent reporter for studying peptide degrons in Escherichia coli. We provide protocols for two different degradation assays. The plate spot assay allows for rapid screening of degron motifs efficiency. It can be used as initial screening method before more thorough evaluation by time-dependent degradation assay. The assays can help to validate the degradation pathway by using single deletion mutant strains which are easily accessible from Keio collection [5]. Alternatively, when protease driving the degradation is not known, the assay can be performed in presence of broad-spectrum protease inhibitor bortezomib. The presented protocols do not require special set up or equipment and can be performed with just fluorescent gel scanner and plate reader which are available in most laboratories, making them a powerful screening strategy before committing to more in-depth degron investigation in vitro [6] described separately in a dedicated chapter.

## 2. Materials

### 2.1. Cloning

1. Inducible plasmid encoding eGFP under arabinose promoter (Addgene plasmid #54762 EGFP-pBAD), *see* **Note 1**.
2. Template-specific primers with overhangs encoding a degron sequence of interest
3. High fidelity DNA polymerase (e.g. Phusion DNA polymerase)
4. 10 mM dNTPs mix
5. Nuclease-free water
6. Thermal cycler
7. DpnI enzyme
8. Digestion buffer
9. DNA loading dye
10. DNA ladder
11. Agarose (molecular biology grade)
12. TAE buffer (Tris base 40 mM, acetic acid 20 mM, EDTA 1 mM)
13. DNA dye (e.g. SYBR™ Safe)
14. Horizontal electrophoresis apparatus and power pack
15. Clean scalpel or gel cutting device
16. Gel-out DNA purification kit or PCR DNA purification kit
17. Nano-spectrophotometer (e.g. Nanodrop)
18. Polynucleotide kinase (PNK)
19. T4 DNA ligase
20. DNA ligase buffer
21. 10 mM ATP
22. Electrocompetent *E. coli* strain compatible with cloning (DH5α or Top10)
23. Thermoblock
24. Luria Bertani (LB)-agar plates with ampicillin (100 µg/mL)
25. Mini-prep DNA isolation kit

### 2.2. Bacterial cultures for degrons expression

1. Electrocompetent *E. coli* strain BW25113 and protease mutant strains available from Keio strain collection [5]
2. Luria Bertani (LB) medium
3. 20% sterile L(+)-arabinose solution
4. Ampicillin 100 mg/mL
5. Kanamycin 15 mg/mL
6. LB-agar plates with 100 µg/mL ampicillin and 0.001 % arabinose or with 100 µg/mL ampicillin, 15 µg/mL kanamycin and 0.001 % L(+)-arabinose
7. Bortezomib stock solutions in DMSO at 50, 25, 10, 5, and 1 mM (optional)
8. Shaking incubator
9. Spectrophotometer with compatible cuvettes

### 2.3. Degron evaluation

1. Ampicillin 100 mg/mL
2. Kanamycin 15 mg/mL
3. Spectinomycin 100 mg/mL
4. M9 medium: (Na_2_HPO_4_ 6 g/L, KH_2_PO_4_ 3 g/L, NaCl 0.5 g/L, NH_4_Cl 1 g/L, MgSO_4_ 120 mg/L, CaCl_2_ 11 mg/L, CaCl_2_ · 2 H_2_O 15 mg/L, Na_2_EDTA · 2 H_2_O 30 mg/L, FeCl_3_ · 6 H_2_O 25 mg/L, CuSO_4_ · 5 H_2_O 2.4 mg/L, MnSO_4_ · H_2_O 1.8 mg/L, ZnSO_4_ · 7 H_2_O 0.27 mg/L, CoCl_2_ 0.27 mg/L, thiamine 10 mg/L, biotin 10 mg/L)
5. 94-well black plate with optic bottom
6. Gel imager with optic filters for GFP (excitation at λ = 488 nm and emission at λ = 520 nm)
7. Microplate reader with shaking and temperature control with optical filters for GFP (excitation at λ = 488 nm and emission at λ = 520 nm)

## 3. Methods

### 3.1. Preparation of plasmids the GFP-degron fusion construct by site-directed mutagenesis

1. Insertion of degron sequences up to 15-20 amino acids can be easily achieved by site-directed mutagenesis. Design the primers according to the orientation of your investigated degrons. If you do not know their orientation, create constructs with both N-terminal and C-terminal fusion to eGFP. When designing your primers, aim for a similar hybridisation temperature (ideally between 60 and 65 °C) for the parts which are complementary to the plasmid template. The plasmid overhangs should encode your degron and should face away from each-other to amplify the entire plasmid while adding the degron in fusion with eGFP (Figure 1). If you wish to investigate longer degron sequences, you might have to consider other cloning techniques (e.g. Gibson assembly).
2. Prepare a 50 µL PCR reaction containing: 10 ng eGFP-pBAD plasmid template 200 µM dNTPs mix 0.5 µM forward primer 0.5 µM reverse primer 1x polymerase specific reaction buffer 2 u high fidelity DNA polymerase Run the reaction in the thermal cycler (*see* **Note 2**) Initial denaturation 98°C 30 s Repeat the following steps 30 times: Denaturation 98°C 10 s Primer annealing 65°C 15 s Elongation 72°C 2 min Final elongation 72°C 5 min Run a 5 µL aliquot of the PCR product on 0.8% agarose gel with DNA dye in 1x TAE buffer to evaluate the presence and purity of your PCR product. Visualise with a gel imager or transilluminator. The template plasmid has a size of 5429 bp and, for short degron peptide motifs, the size difference between modified and parental plasmid might not be distinguishable on the gel.
3. To remove the template DNA from the reaction mix, dilute the PCR product to 89 µL (so that the final reaction volume would be 100 µL) with nuclease-free water, add the enzyme-specific buffer to the final concentration of 1x, and DpnI enzyme. Incubate the reaction at 37°C for 15-30 min, according to the manufacturer’s instructions. Purify the DNA (*see* **Note 3**). Quantify the DNA concentration with a nano-spectrophotometer.
4. Assemble the circularisation reaction with PNK to phosphorylate 5’ DNA ends and T4 ligase in a total volume of 20 µL: 100 ng purified DNA 1x DNA ligase buffer 250 µM ATP (optional) 5 u PNK 2.5 u T4 DNA ligase Incubate the reaction for 1 h at room temperature. Alternatively, the reaction might be left overnight at 4°C.
5. Transform 5 µL of the ligation mixture to a 50 µL aliquot of electrocompetent cloning-compatible *E. coli* strain. Grow bacteria on selection LB-agar plates with ampicillin at 37°C overnight. The following day inoculate a couple of colonies in 5 mL of LB with 100 µg/mL ampicillin and grow for 16-20 h at 37°C in a shaking incubator. Isolate plasmid DNA and confirm the insertion of the degron sequence by DNA sequencing.

**Figure 1.**
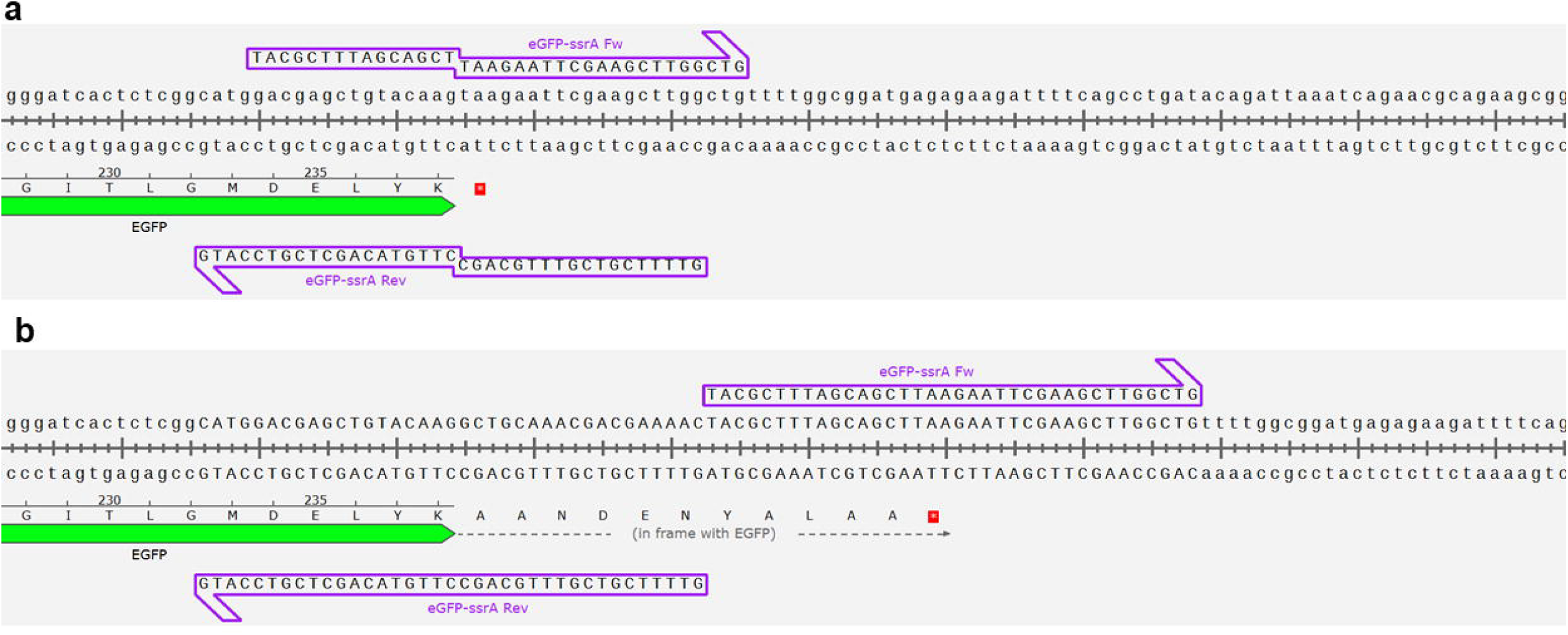
An example of primer design for introducing ssrA degron at the C-terminus of eGFP in EGFP-pBAD plasmid. Primers are outlined with purple arrows. (**a**) A representation of the primer hybridisation sites in the parental plasmid. Parts of forward and reverse primers hybridise to the vector and the overhangs encode ssrA degron. (**b**) A representation of the same plasmid region after introducing the ssrA degron by site-directed mutagenesis

### 3.2. Preparation of bacterial cultures for degron evaluation

Transform plasmids encoding eGFP-degron fusions in *E. coli* strain BW25113. Include control with unmodified degron-less eGFP construct and bacteria transformed with an empty vector (such as pBAD-6xHis-TEV, *see* **Note 4**). If you know the protease recognising the investigated degron motifs, you can transform the plasmids in parallel to a single gene mutant strain (e. g. from Keio collection). Grow the bacteria overnight at 37°C on plates with 100 µg/mL ampicillin (wild type BW25113) or 100 µg/mL ampicillin and 15 µg/mL kanamycin (mutant strains). Inoculate single colonies of transformed bacteria in 5 mL LB with appropriate selection antibiotics and grow overnight at 37°C in a shaker incubator. At this point, you can use aliquots of the cultures for a spot assay on agar plates (see below). For the dynamic degradation evaluation in liquid cultures, dilute the cultures 1:100 in 5 mL of LB with selection antibiotics and grow to mid-exponential phase (OD_600_ = 0.5-0.6). Induce the cultures with 0.005% L-arabinose and incubate overnight at 18°C with shaking.

### 3.3. Plate spot assay

Dilute the overnight cultures 1:100 in sterile LB. Place 3 µL drops on LB agar plates with selection antibiotics (100 µg/mL ampicillin for BW25113 strain and 100 µg/mL ampicillin with 15 µg/mL kanamycin for Keio mutant strains) supplemented with 0.001% L-arabinose. Incubate the plates overnight at 37°C. Image plates on a gel imager with optic filters for GFP or similar (excitation at λ = 488 nm and emission at λ = 520 nm, such as Alexa Fluor 488 or fluorescein) (Figure 2).

**Figure 2.**
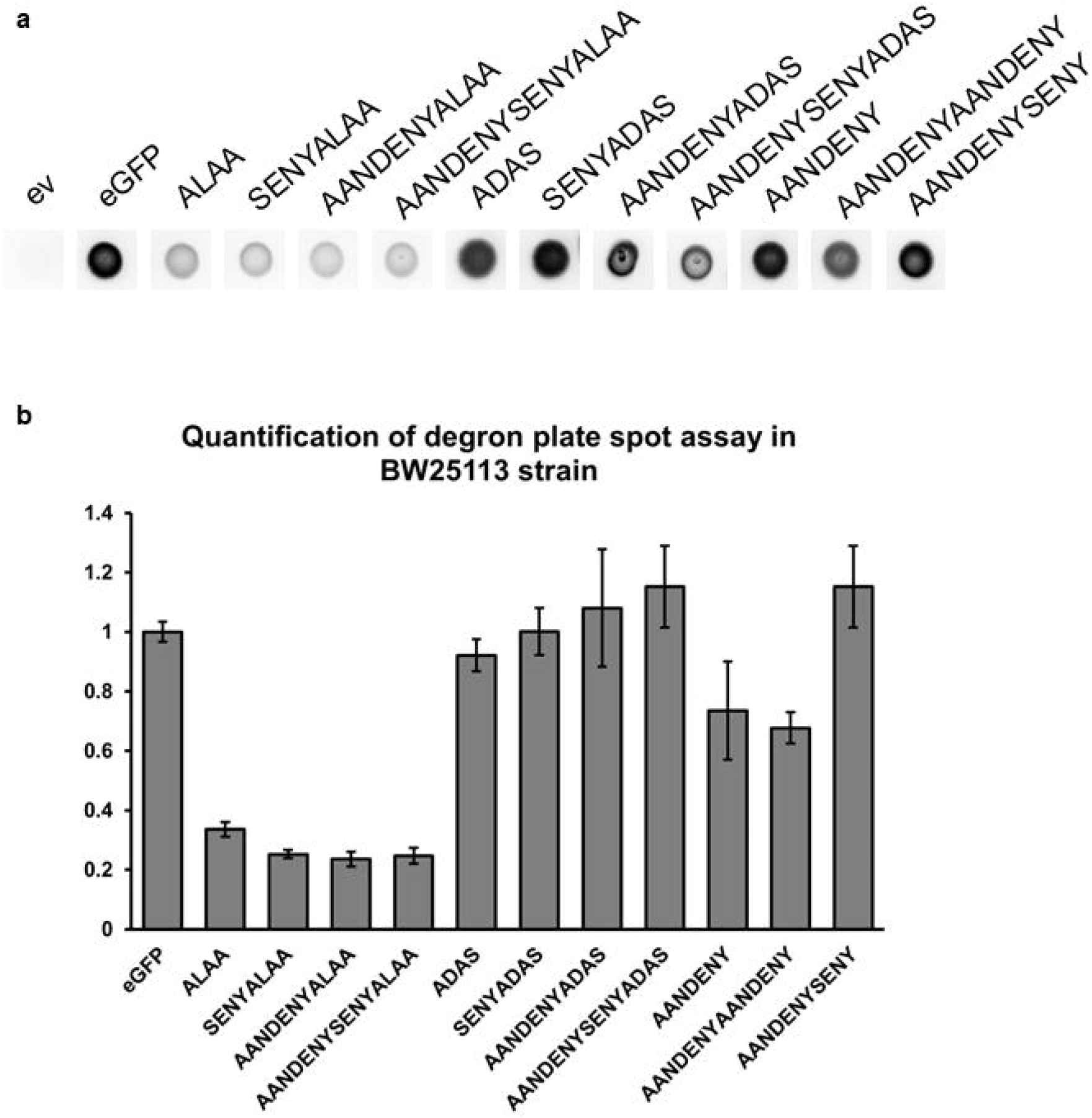
An example of plate spot assay results in BW25113 strain. (**a**) Pictures of bacterial spot colonies grown on LB-agar in presence of 0.005% L(+)-arabinose expressing eGFP in fusion with peptide degrons. (**b**) Quantification of colony fluorescence. The bars represent mean values from one biological repeat performed in triplicate. The error bars represent SEM.

### 3.4. Time-dependent microplate degradation assay

Prepare a 96-well black plate with optic bottom. Assemble the plate on ice to reduce the degradation of eGFP-degron constructs during preparation of the plate. Pipette 190 µL of M9 medium with 100 µg/mL ampicillin (or 100 µg/mL ampicillin and 15 µg/mL kanamycin for measurements in mutant strains), 100 µg/mL spectinomycin, and 0.2% glucose to ensure the expression arrest of the constructs. Then add 10 µL of the induced cultures to the wells (to achieve 1:20 dilution) in triplicates. Place the plate in a microplate reader pre-heated to 30°C and measure the fluorescence and OD_600_ in the wells at 15 minutes interval for 6 hours with shaking in between the measurements, *see* **Note 5** (Figure 3 a-c).

**Figure 3.**
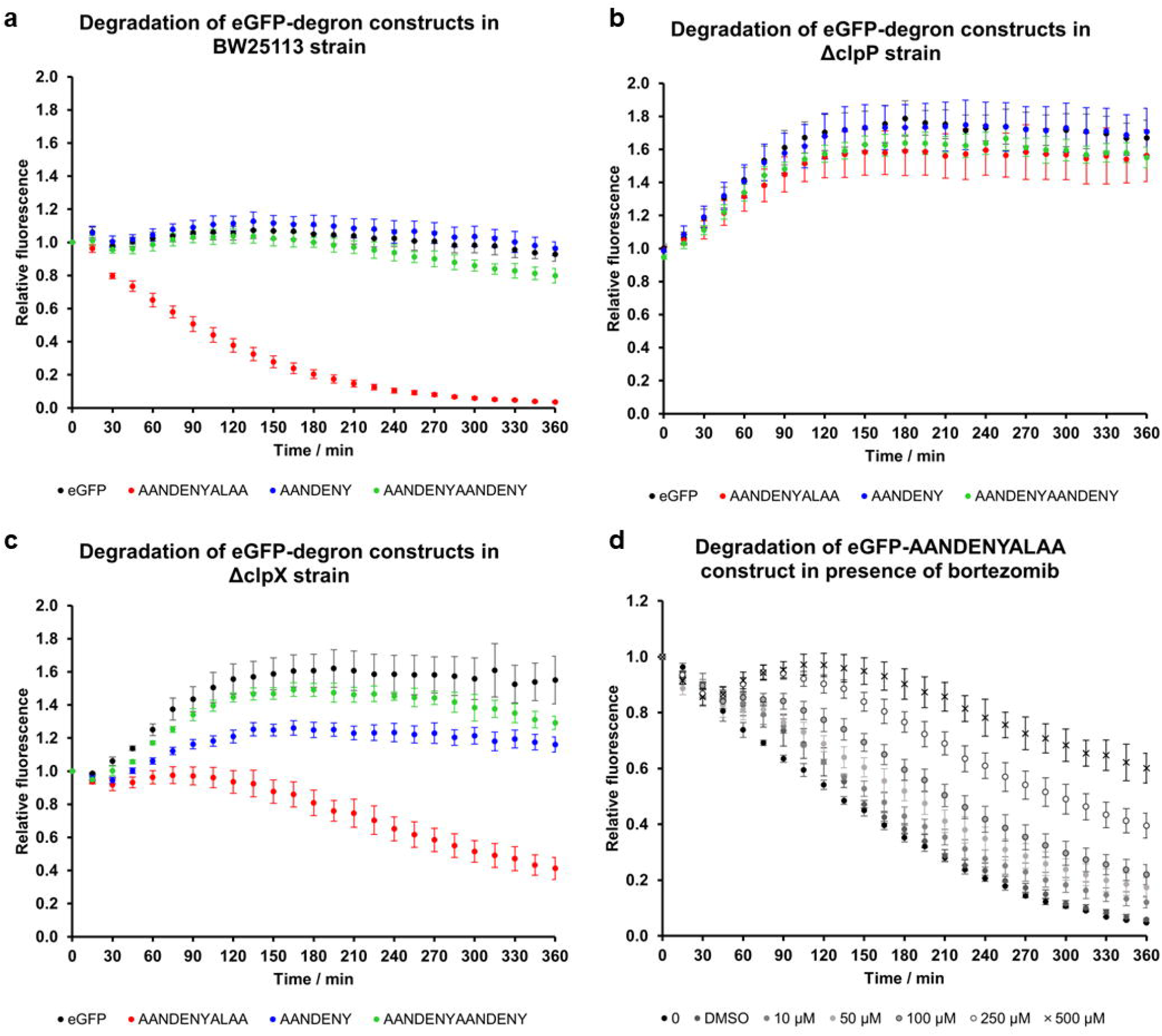
Degradation of eGFP-degron constructs measured in liquid cultures in a 96-well plate format, the plotted values represent mean of 3 biological replicates and the error bars represent SEM (**a**) degradation of selected eGFP-degron constructs in wild type BW25113 strain, (**b**) degradation of eGFP-degron constructs in protease mutant strain ΔclpP, (**c**) degradation of eGFP-degron constructs in unfoldase mutant strain ΔclpX, (**d**) degradation of chosen eGFP-AANDENYALAA (ssrA) constructs in wild type BW25113 strain in presence of increasing concentration of bortezomib.

### 3.5. Bortezomib inhibition assay

For bortezomib inhibition assay prepare the bacterial cultures as described above. When preparing the 96-well plate with M9 medium, add bortezomib in DMSO at concentrations 500, 250, 100, 50, 10 µM or 1% DMSO. Then, dilute the cultures 1:20 in the prepared microplate and measure in a microplate reader as described above (Figure 3 d).

### 3.6. Data evaluation

Plate spot test can be quantified in image quantification software such as Fiji [7]. Select the region of interest (ROI) by drawing circles around the bacterial spots. Quantify the signal intensity by measuring brightness of the spots. To account for the background, subtract the signal of bacteria expressing the empty vector.

To evaluate the results from plate reader measurements, normalise the fluorescence value of each datapoint by dividing the fluorescence value by OD_600_ value. Subtract the average value obtained for bacteria expressing empty vector from data points at each timepoint. Then, normalise the datapoints to 100% at 0 min time point for each evaluated degron. Plot the processed data (normalised fluorescence vs. time) and determine t_1/2_ of each eGFP-degron construct (*see* **Note 6**).

## 4. Notes

1. The protocol can be adapted to other plasmids, provided the chosen expression strain is compatible with the promoter (e.g. T7 promoter with *E. coli* strains DE3 expressing T7 RNA polymerase). Arabinose promoter of pBAD plasmid provides a tight expression control [8] and is compatible with expression in strains from the Keio collection [5] which provides an easy access to a wide variety of single-gene deletion mutants for validation of degron-specific proteases or adaptor proteins.
2. The described conditions were optimised for Phusion DNA polymerase. Other polymerases might require adjusting the temperature and time of the different PCR stages. If initial conditions do not yield enough of the DNA product or you observe a significant amount of non-specific product amplification, you might have to test other annealing temperatures to optimise the reaction conditions or pool several 50 µL reactions.
3. Depending on the purity of your DNA product you might directly purify the DNA with a clean-up kit. If you observe the formation of non-specific DNA products, run the entire volume of the reaction on agarose gel (as above). Excise the band with the DNA product of expected length using a clean scalpel or a gel-cutting device and extract the DNA with a DNA gel-out kit.
4. An empty pBAD vector can be obtained by deletion of eGFP by site-directed mutagenesis using analogous protocol as described in **3.1**. The primers for deletion of the eGFP should be complementary to the parts flanking the DNA sequence encoding the eGFP.
5. The kinetic interval and measurement time can be adjusted according to the degradation speed of specific degrons, however evaporation of the sample will be more significant for longer plate incubations.
6. Plots can be analysed using different data analysis software such as GraphPad Prism.

